# The zinc transporter ZnuABC is critical for the virulence of *Chromobacterium violaceum* and contributes to diverse zinc-dependent physiological processes

**DOI:** 10.1101/2021.06.03.447022

**Authors:** Renato E. R. S. Santos, Waldir P. da Silva Júnior, Simone Harrison, Eric P. Skaar, Walter J. Chazin, José F. da Silva Neto

## Abstract

*Chromobacterium violaceum* is a ubiquitous environmental bacterium that causes sporadic life-threatening infections in humans. How *C. violaceum* acquires zinc to colonize environmental and host niches is unknown. In this work, we demonstrated that *C. violaceum* employs the zinc uptake system ZnuABC to overcome zinc limitation in the host, ensuring the zinc supply for several physiological demands. Our data indicated that the *C. violaceum* ZnuABC transporter is encoded in a *zur*-CV_RS15045-CV_RS15040-*znuCBA* operon. This operon was repressed by the zinc uptake regulator Zur and derepressed in the presence of the host protein calprotectin (CP) and the synthetic metal chelator EDTA. A Δ*znuCBA* mutant strain showed impaired growth under these zinc-chelated conditions. Moreover, the deletion of *znuCBA* provoked a reduction in violacein production, swimming motility, biofilm formation, and bacterial competition. Remarkably, the Δ*znuCBA* mutant strain was highly attenuated for virulence in an *in vivo* mouse infection model and showed a low capacity to colonize the liver, grow in the presence of CP, and resist neutrophil killing. Overall, our findings demonstrate that ZnuABC is essential for *C. violaceum* virulence, contributing to subvert the zinc-based host nutritional immunity.

## INTRODUCTION

The transition metal zinc plays an essential role in both eukaryotic and prokaryotic cells (1, 2). Zinc functions as a structural and catalytic cofactor of several classes of proteins such as proteases (3), endopeptidases (4, 5), metallo-β-lactamases (6), and transcription factors (2). Large fractions of the proteome of bacteria (about 5-6%) and vertebrates (about 8-10%) are zinc-binding proteins (2, 7). Maintenance of bacterial zinc homeostasis requires the coordinated expression of transporters for zinc acquisition and efflux (8). In most Gram-negative bacteria, the Zinc Uptake Regulator Zur is the main metal-sensing regulator that represses genes encoding zinc acquisition systems, such as the ATP-binding cassette (ABC) transporter ZnuABC (8–10). The ZnuABC is a high-affinity tripartite zinc importer composed of a soluble periplasmic zinc-binding protein (ZnuA), a transmembrane protein (ZnuB), and an ATPase (ZnuC) (8, 10). In some cases, Zur also acts as an activator of zinc efflux systems under zinc excess, such as in *Streptomyces coelicolor* (9), *Xanthomonas campestris* (11), and *Caulobacter crescentus* (12).

Zinc also plays a central role during bacterial infection (1, 13). Vertebrate hosts use distinct strategies to make zinc and other metals unavailable to bacterial pathogens, a process termed nutritional immunity (13). During infection, neutrophils express a large amount of calprotectin (CP), a S100A8/S100A9 heterodimer of the S100 family of EF-hand calcium-binding proteins (14, 15). CP binds zinc, manganese, and ferrous iron with high affinity (16, 17) and causes growth inhibition of a large variety of bacteria (18–22). On the other hand, bacteria overcome CP-mediated zinc chelation by expressing high affinity zinc transporters, such as the ZnuABC, as demonstrated in *Salmonella enterica* serovar Typhimurium (19) and *Acinetobacter baumannii* (23, 24). The battle between hosts and bacteria also involves zinc intoxication. For instance, macrophages deliver zinc-containing vesicles to promote microbial clearance, while bacteria express zinc efflux systems to alleviate zinc toxicity (25, 26).

Metal homeostasis is critical for environmental pathogenic bacteria, which need to deal with metal fluctuations in free-living and host-associated conditions. In previous works, we demonstrated the importance of iron for the virulence and the physiology of *Chromobacterium violaceum* (27, 28), a Gram-negative environmental opportunistic pathogen. *C. violaceum* causes human infections with mortality rates ranging from 35 to 53% of the cases (29, 30). Several traits contribute to *C. violaceum* pathogenicity, such as the high intrinsic antibiotic resistance (29, 30), the secretion of effectors by a Cpi1 type III secretion system (31–33), and the production of catecholate-type siderophores regulated by the Ferric Uptake Regulator Fur (27, 28). However, the relevance of zinc homeostasis for the physiology and virulence of *C. violaceum* and how this bacterium acquires zinc are largely unknown.

In this work, we characterized the high affinity ABC-type transporter ZnuABC of *C. violaceum*. We demonstrated that ZnuABC is encoded in a Zur-repressed *zur*-CV_RS15045-CV_RS15040-*znuCBA* operon. A Δ*znuCBA* mutant strain showed impaired growth under zinc-chelated conditions and reduced violacein production, swimming motility, biofilm formation, and bacterial competition. Using growth curves under CP-treatment, neutrophil killing assays, and an *in vivo* mouse infection model, we demonstrated that ZnuABC exerts a key role in *C. violaceum* virulence by overcoming the host-imposed and CP-mediated zinc limitation.

## RESULTS

### The *C. violaceum* ZnuABC transporter is encoded in a Zur-repressed *zur*-CV_RS15045-CV_RS15040-*znuCBA* operon

*C. violaceum* possesses two members of the Ferric Uptake Regulator (Fur) family of transcription factors, the previous characterized Fur (27) and the undescribed Zinc Uptake Regulator Zur (CV_RS15050). Genome inspection revealed that *zur* composes a putative operon with other five genes (Fig. 1a): CV_RS15045, encoding an uncharacterized protein belonging to the Cluster of Orthologous Groups (COG) COG0523, CV_RS15040, encoding a small accessory hypothetical protein, and the *znuC*, *znuB*, and *znuA* genes, which encode a putative ZnuABC transport system (Fig. 1a). Zur represses Zur-regulated genes by binding in a palindromic regulatory sequence, known as Zur-box (9–12). To identify a putative Zur binding site in *C. violaceum*, we searched for conserved DNA motifs upstream of *zur* in several genomes of *Chromobacterium*, using the motif discovery tool MEME (34). This analysis returned a possible Zur-binding motif (Fig. 1b) located upstream of *zur* that resembles the previously characterized Zur-boxes of *Pseudomonas protegens* (35) and *Neisseria meningitidis* (36). We designed primers in three genes of the possible operon (Fig. 1a) to verify by reverse transcriptase-polymerase chain reaction (RT-PCR) if they were co-transcribed. Firstly, we confirmed the absence of DNA contamination in the isolated total RNA by PCR (Fig. 1c, upper panel). Using the same combination of primers, we detected co-transcription by RT-PCR (Fig. 1c, down panel), indicating the occurrence of a *zur*-CV_RS15045-CV_RS15040-*znuCBA* operon in *C. violaceum*. To investigate Zur-mediated regulation of this operon in *C. violaceum*, we generated a fusion of the promoter region of *zur* (p*zur*) with a promoterless *lacZ* gene. The β-galactosidase activity assays revealed that in the wild-type (WT) strain, the expression of this operon doubled from zinc-replete (LB and LB plus zinc) to zinc-chelated conditions (EDTA and TPEN treatments) (Fig. 1d). As a control, the p*zur*-*lacZ* fusion was unresponsive to iron-replete (FeSO_4_) or iron-chelated conditions (2,2’-DP treatment) (Fig. 1d). In the Δ*zur* strain, regardless of the treatment, the p*zur* expression was derepressed, demonstrating that Zur acts as a repressor of this operon (Fig. 1d). Altogether, these results indicated that *C. violaceum* possesses a zinc-responsive, Zur-regulated operon containing six genes potentially involved in zinc homeostasis. Hereafter, we will focus on the characterization of the *C. violaceum* ZnuABC.

**Fig 1.**
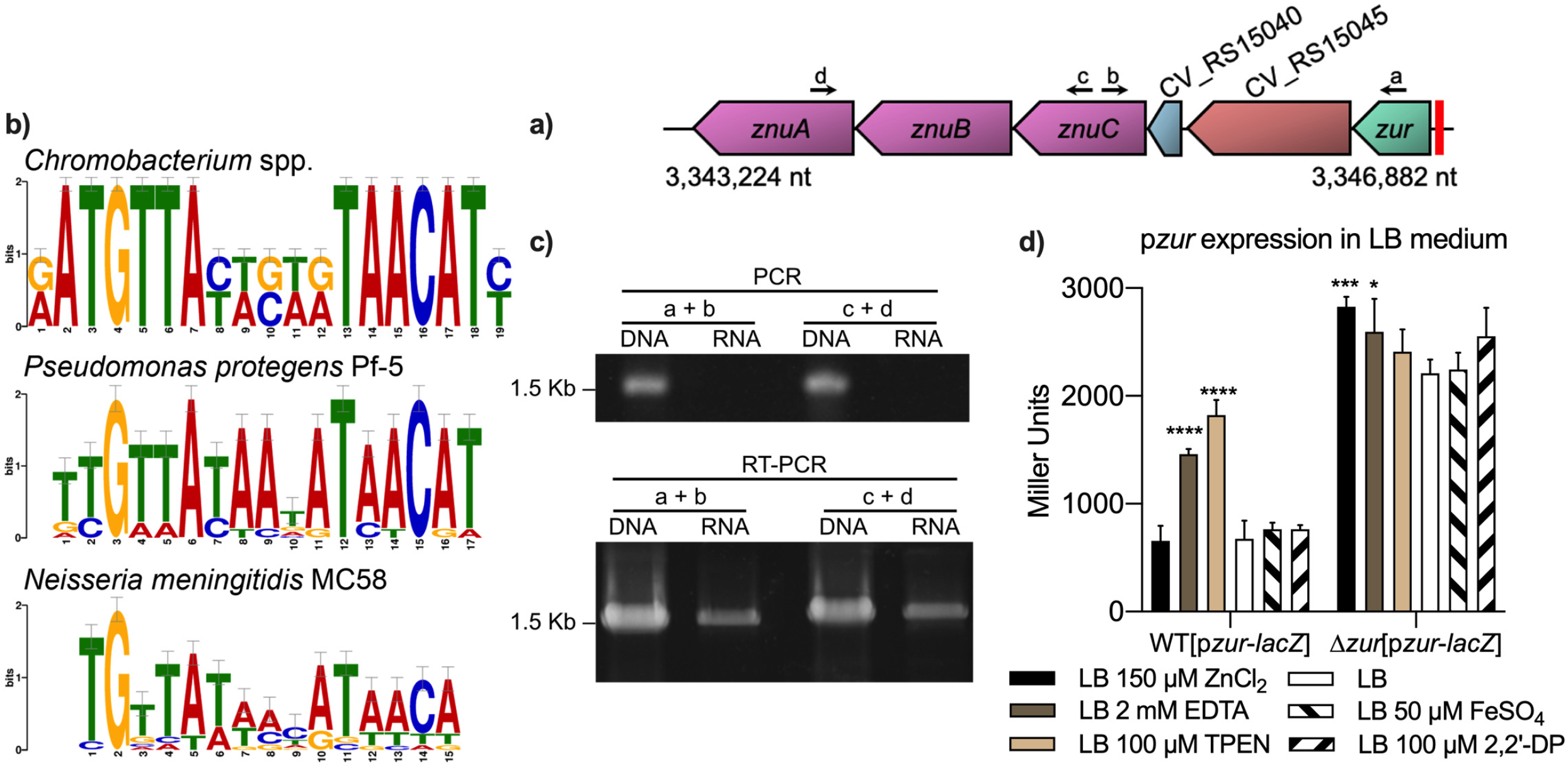
The *C. violaceum* ZnuABC is encoded in a Zur-repressed, zinc responsive operon. (A) Genomic context of the genes *zur*, CV_RS15045, CV_RS15040, and *znuCBA*. The red box upstream of *zur* indicates a predicted Zur-box. The arrows indicate the annealing positions of the primers used in the RT-PCR reactions. (B) The putative Zur binding site predicted upstream of *zur* from *Chromobacterium* spp. This DNA binding motif matches the known Zur-boxes of *Pseudomonas protegens* Pf-5 and *Neisseria meningitidis* MC58. (C) Co-transcription of genes of the *zur*-CV_RS15045-CV_RS15040-*znuCBA* operon. Control PCR reactions (upper panel) using genomic DNA or RNA as templates. RT-PCR reactions (down panel) using the indicated primers (“a” and “b”, 1552 bp; “c” and “d”, 1613 bp) confirmed co-transcription of the genes of the operon. (D) Expression of the promoter region of *zur* (p*zur*-*lacZ*) determined by β-galactosidase activity assay. Strains harboring the p*zur*-*lacZ* fusion were grown in LB medium, and the cultures were treated for 2 h with the indicated concentrations of ZnCl_2_, EDTA, TPEN, FeSO_4_, and 2,2’-DP. These experiments were performed in three independent biological replicates. Statistical analysis performed by One-way ANOVA followed by Dunnett’s multiple comparison test. Samples were compared to the untreated LB culture of WT[p*zur*-*lacZ*] or Δ*zur*[p*zur*-*lacZ*]. \**P*=0.0358, \*\*\**P*=0.0008, \*\*\*\**P*<0.0001, where not indicated = not significant.

### The absence of ZnuABC impairs the growth of *C. violaceum* in zinc limitation

To study how *C. violaceum* copes with zinc availability *in vitro*, we determined the minimal inhibitory concentration (MIC) of the zinc-chelating agent EDTA. The WT and Δ*znuCBA* strains showed the same MIC for EDTA (128 mM) (Fig. 2a). Considering that EDTA can chelate other divalent cations, we tested different combinations of metals (zinc or iron) and metal-chelators (EDTA or 2,2’-DP) to assess the growth after 24 h cultivation (Fig. 2b) and the growth inhibition by disk diffusion assays (Fig. 2c, d). Under a condition of EDTA plus zinc (LB 1 mM EDTA 250 μM ZnCl_2_), the Δ*znuCBA* mutant had an impaired growth compared to that of the WT and complement strains (Fig. 2b). In agreement, in the disk diffusion assays, the Δ*znuCBA* mutant was more sensitive to EDTA treatment (inhibition halos from 62.5 mM EDTA) than the WT or the complemented strain (inhibition halos from 250 mM EDTA) (Fig. 2c). As a control, all strains showed the same profile of growth (Fig. 2b) or growth inhibition (Fig. 2d) under iron chelated (2,2’-DP) or iron supplemented conditions (FeSO_4_). These data demonstrate that ZnuABC is required for growth under zinc, but not iron, limitation. To further characterize the effects of zinc deficiency in a Δ*znuCBA* mutant strain, we performed growth curves in LB medium with different zinc conditions. The Δ*zur* and Δ*znuCBA* mutant strains showed the same growth profile as that of the WT strain in LB (Fig. 3a) or LB plus 1 mM EDTA (Fig. 3b), raising the possibility that the intracellular zinc content would still be sufficient to maintain proper cell physiology. To investigate this hypothesis, we treated all pre-inocula with 1 mM EDTA and measured growth curves in LB (Fig. 3c) and LB plus 1 mM EDTA (Fig. 3d). In these experiments, the growth of Δ*znuCBA* was mildly compromised in LB (Fig. 3c) and severely compromised in LB plus 1 mM EDTA (Fig. 3d) compared to that of the WT strain. In both cases, these phenotypes were reverted in the complemented strain. Zinc supplementation in the growth curves using EDTA-treated pre-inocula was not able to fully restore the growth defect of the Δ*znuCBA* mutant strain (Fig. 3e). Overall, these experiments demonstrate that the ZnuABC transporter plays an important role in zinc acquisition in *C. violaceum*, especially under severe zinc scarcity, when the intracellular zinc pool was previously depleted.

**Fig 2.**
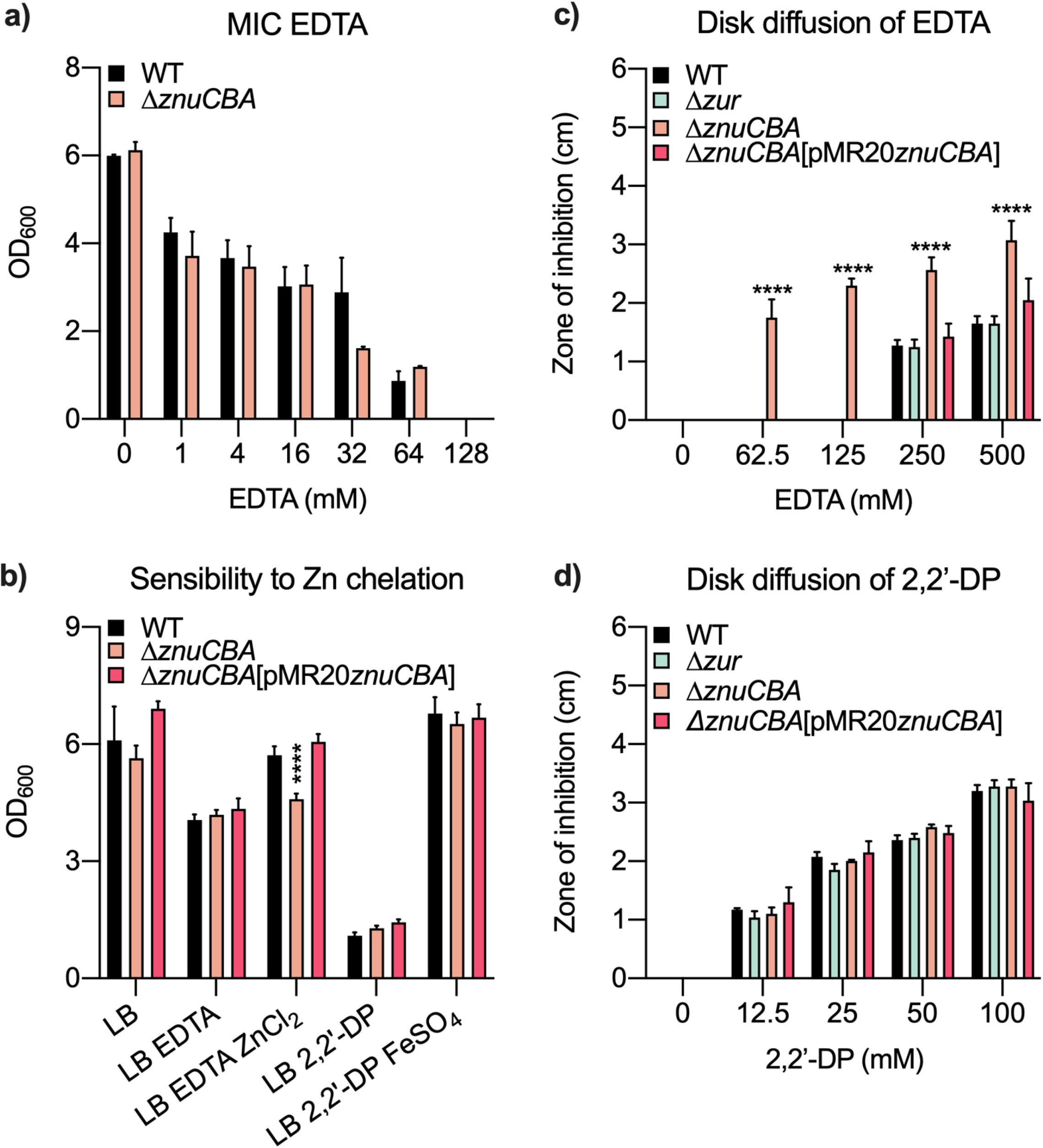
The absence of ZnuABC affects the sensitivity of *C. violaceum* to zinc, but not to iron, limitation. (A) The MIC of EDTA in LB medium was the same for the WT and *ΔznuCBA* strains. Data from three biological replicates are shown as mean and standard deviation. Statistical analysis performed by unpaired *t-*test for each EDTA concentration indicated the absence of statistical significance. (B) Growth of the indicated strains in LB in different conditions of zinc (EDTA, ZnCl_2_) and iron (2,2’-DP and FeSO_4_) availability. The growth was evaluated after 24 h cultivation. Statistical analysis performed by One-way ANOVA followed by Dunnett’s multiple comparison test. Samples were compared to the WT strain in each condition. \*\*\*\**P*<0.0001, where not indicated = not significant. (C and D) Disk diffusion assays for the chelating agents EDTA (C) and 2,2’-DP (D) with the indicated strains. The diameter of the zone of inhibition was measured in three independent biological replicates. Statistical analysis performed by One-way ANOVA followed by Dunnett’s multiple comparison test. Samples were compared to the WT strain in each concentration of EDTA or 2,2’-DP. \*\*\*\**P*<0.0001, where not indicated = not significant.

**Fig 3.**
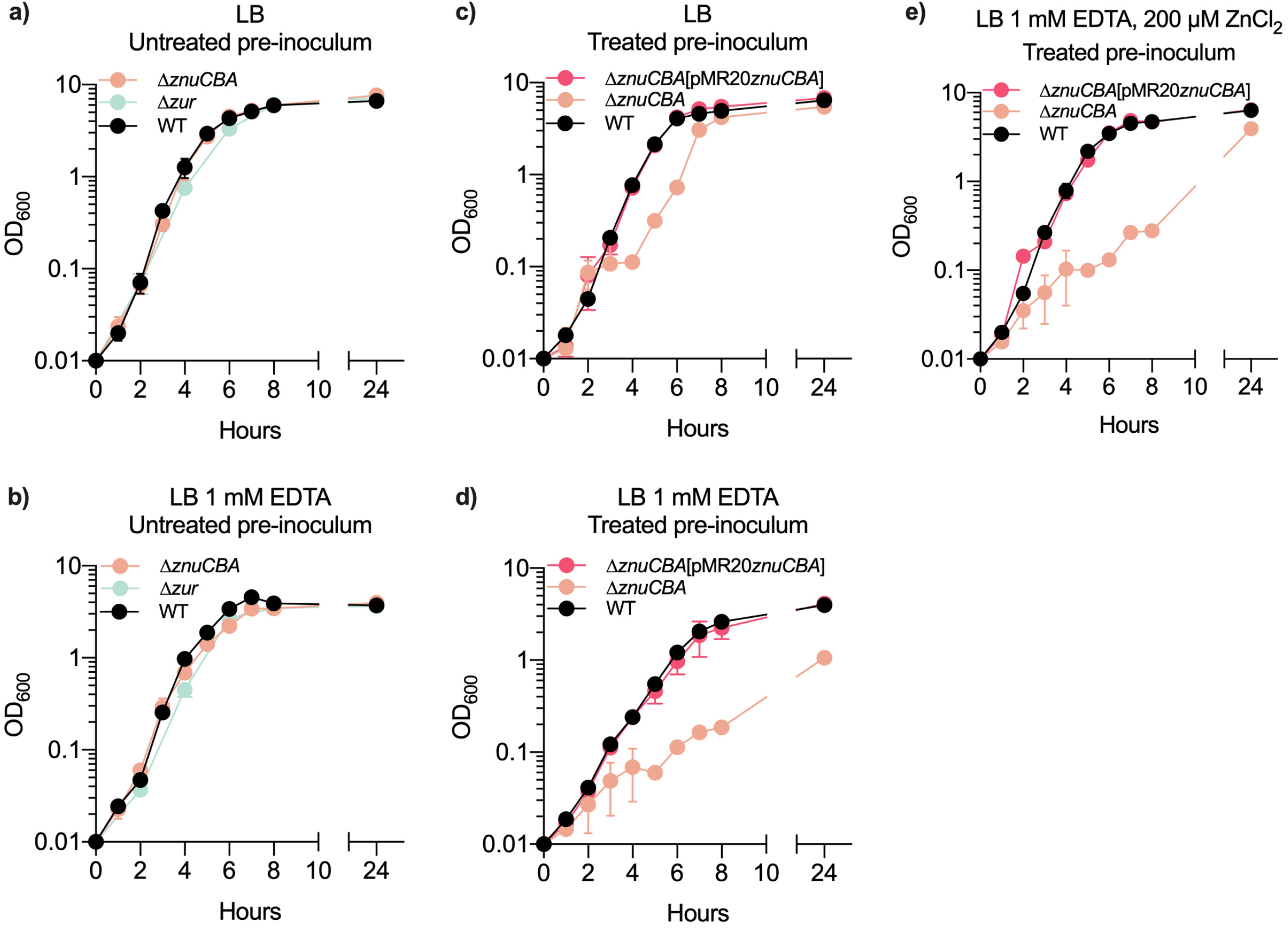
Intracellular zinc depletion by EDTA treatment impairs the growth of Δ*znuCBA*. Growth curves of the indicated strains in LB (A and C) and LB supplemented with 1 mM EDTA (B and D), with pre-inocula untreated (A and B) or treated with 1 mM EDTA (C and D). (E) Growth curves of EDTA-treated pre-inocula in LB supplemented with 1 mM EDTA and 200 μM ZnCl_2_. Growth curves were determined in three independent biological replicates and are represented as the mean with standard deviation.

### *C. violaceum* ZnuABC is required for growth in zinc limitation imposed by calprotectin

Calprotectin (CP) is a zinc, manganese, and iron chelating protein released from neutrophils that plays a central role in host defense against bacterial pathogens (17–19, 21). To investigate the antimicrobial role of CP against *C. violaceum*, we used a wild-type CP (CP WT) and a selectively inactivated CP with mutations in both the noncanonical (SI) and the canonical (SII) transition metal-binding sites (CP ΔSI/SII) (17). Determination of the 50% growth inhibitory concentration (IC_50_) of CP WT indicated that *C. violaceum* is more resistant to CP than an *Escherichia coli* DH5α strain, used as a control (IC_50_ of 534.6 ± 13.8 and 287.5 ± 22.3, respectively). We measured growth curves in LB supplemented with 500 μg/ml of CP WT or CP ΔSI/SII, using CP treated or untreated pre-inocula (Fig. 4). When bacteria were treated with CP WT and the growth curve was performed in LB with CP WT, the Δ*znuCBA* mutant strain showed impaired growth compared to that of the WT and complement strains (Fig. 4a). This effect was not detected when CP ΔSI/SII was present in both the pre-inocula and inocula (Fig. 4b), indicating a metal binding-dependent CP effect. When CP WT was only present in the growth curve, without pre-treatment, all strains presented the same growth profile (Fig. 4c), confirming the requirement of ZnuABC for growth when the intracellular zinc pool was depleted. We also measured the β-galactosidase activity of the p*zur*-*lacZ* fusion in the WT strain in the presence of CP (Fig. 4d). The expression of p*zur* increased two-fold with the CP WT or EDTA treatments compared to the untreated control. Treatment with the CP ΔSI/SII had no significant effect (Fig. 4d). These data indicate that CP imposes zinc limitation to *C. violaceum*, which in turn responds by expressing ZnuABC to overcome CP-mediated zinc chelation.

**Fig 4.**
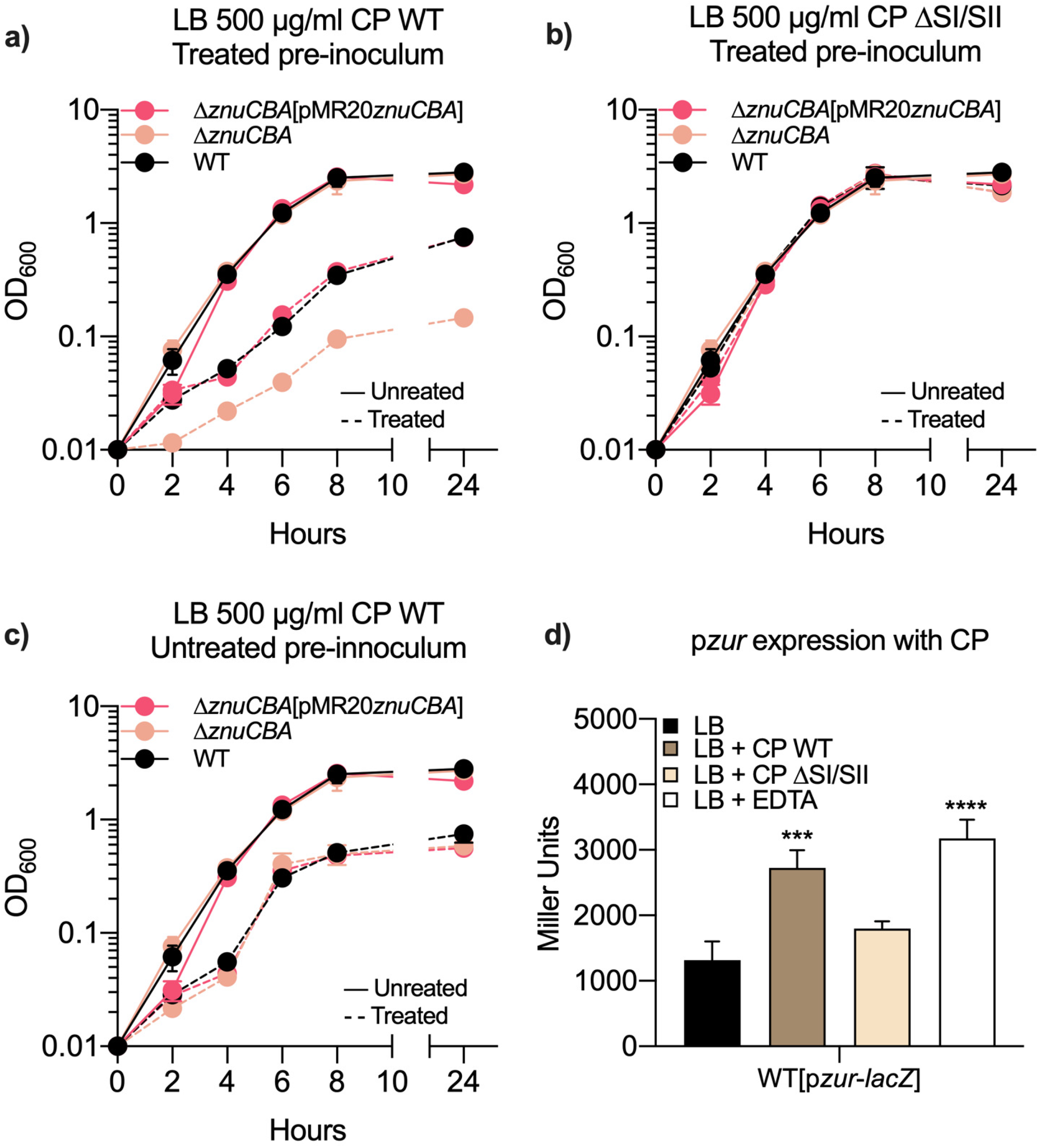
Calprotectin decreases the growth of Δ *znuCBA*. (A-C) Growth curves of the indicated strains determined in the presence of 500 μM CP WT (A and C) or 500 μM CP ΔSI/SII (B). (A) Pre-inocula treated with 500 μg/ml CP WT and growth curves performed without (continuous lines) or with (dashed lines) 500 μg/ml CP WT. (B) Pre-inocula treated with 500 μg/ml CP ΔSI/SII and growth curves performed without (continuous lines) or with (dashed lines) 500 μg/ml CP ΔSI/SII. (C) Untreated pre-inocula used in growth curves performed without (continuous lines) or with (dashed lines) 500 μg/ml CP WT. (D) Effect of CP on the expression of the promoter region of *zur* (p*zur*-*lacZ*). The β-galactosidase activity assays were performed 4 h after the treatment with 500 μM CP WT, 500 μM CP ΔSI/SII, or 1 mM EDTA. Experiments were performed in three independent biological replicates and are represented as the mean with standard deviation. Statistical analysis performed by One-way ANOVA followed by Dunnett’s multiple comparison test. Samples of each treated condition were compared with the samples in LB. \*\*\**P*=0.0004, \*\*\*\**P*<0.0001.

### ZnuABC ensures the supply of zinc required for violacein production, swimming motility, biofilm formation, and bacterial competition

During the MIC assays (Fig. 2a), we observed that EDTA had an inhibitory effect on the production of the pigment violacein. Further characterization indicated that the production of violacein was inhibited at 1 mM EDTA in the WT strain, whereas in the Δ*znuCBA* mutant, inhibition was achieved from 0.031 mM EDTA. This phenotype was reverted with the addition of zinc and after genetic complementation (Fig. 5a). These data suggest that zinc is required for violacein production in *C. violaceum*. We tested whether other phenotypic traits, such as swimming motility and biofilm, could be affected in zinc limitation on the dependence of ZnuABC. Motility tests in semi-solid M9CH agar plates indicated that the flagellar swimming motility of *C. violaceum* WT decreased upon the addition of EDTA. Still, when zinc was added together with EDTA, the motility matched that of the untreated M9CH condition (Fig. 5b). The Δ*znuCBA* mutant strain was less motile than the WT, Δ*zur*, and Δ*znuCBA*[pMR20*znuCBA*] strains (Fig. 5c). These results indicate that *C. violaceum* requires zinc acquisition via ZnuABC for proper swimming motility. We quantified biofilm formation on static conditions of cultures grown in glass tubes in LB or LB plus 100 μM EDTA (Fig. 5d). The biofilm formation in LB was not affected in the Δ*zur* or Δ*znuCBA* mutant strains compared to that of the WT strain.

**Fig 5.**
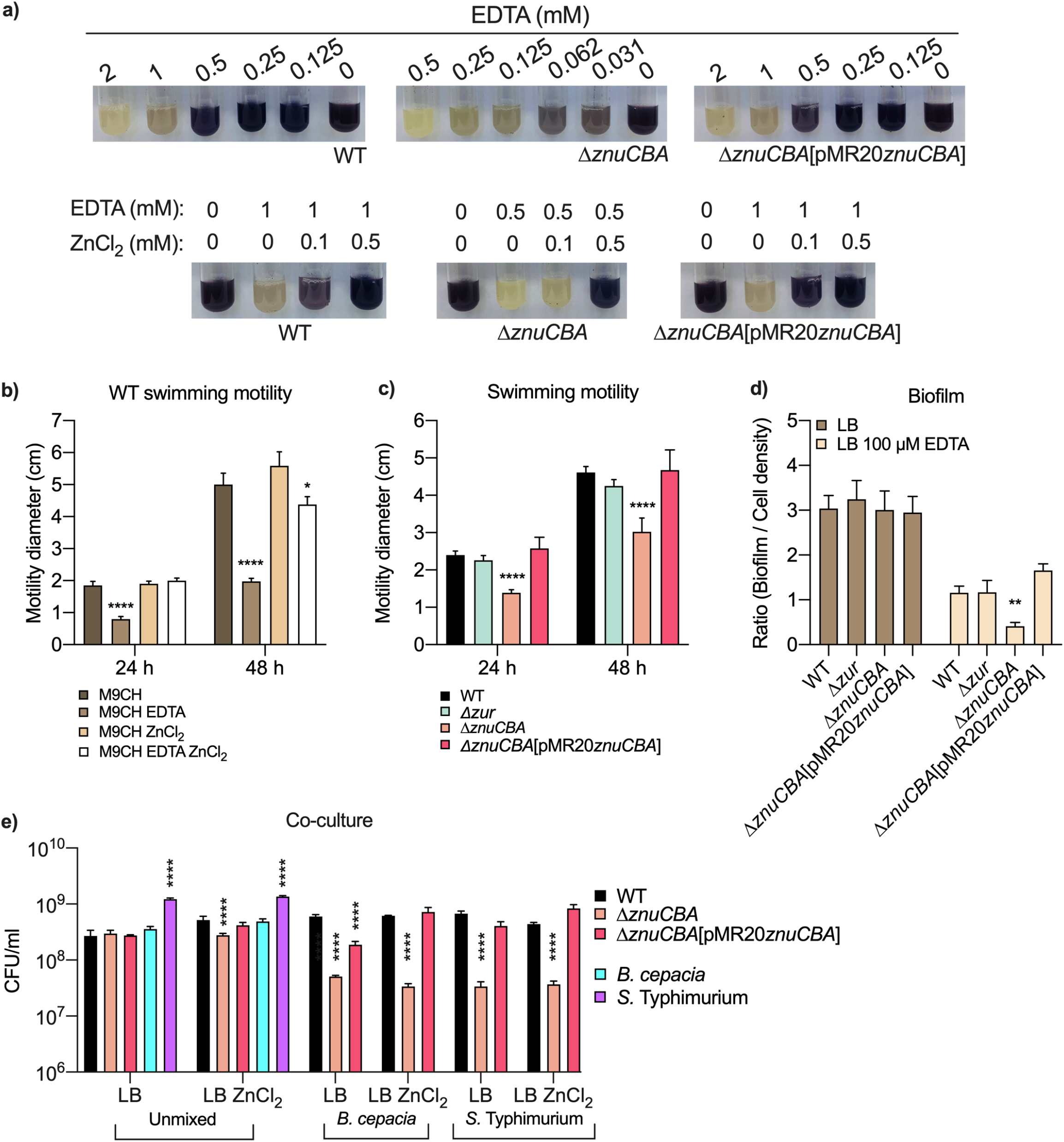
Zinc limitation or mutation of *znuCBA* decrease violacein production, swimming motility, biofilm formation, and bacterial competition. (A) Effect of EDTA and zinc on violacein production in the indicated strains. Photos acquired after 24 h cultivation. (B and C) Effect of EDTA and zinc on swimming motility. (B) WT strain in M9CH or M9CH plus 1 mM EDTA, 200 μM ZnCl_2_, or both EDTA and ZnCl_2_. (C) The indicated strains in M9CH. The halo diameter of bacterial spreading was measured at 24 h and 48 h. Experiments were performed in three independent biological replicates and are represented as the mean with standard deviation. Statistical analysis performed by One-way ANOVA followed by Dunnett’s multiple comparison test. Comparisons were halo diameter of each treated condition with untreated M9CH (A) or mutant strains with WT strain (B). \**P=*0.039, \*\*\*\**P*<0.0001. (D) Effect of EDTA on biofilm in the indicated strains. Static biofilm was determined by crystal violet staining after 24 h of cultivation in LB or LB plus 100 μM EDTA. Data were plotted as the ratio of biofilm quantification by cell density of the cultures (OD_600_). Statistical analysis performed by One-way ANOVA followed by Dunnett’s multiple comparison test. \*\**P*=0.003. (E) The absence of ZnuABC affects *C. violaceum* competition. Interspecies competition of the indicated *C. violaceum* strains co-cultured with *Burkholderia cepacia* ATCC 17759 or *Salmonella* Typhimurium ATCC 14028. Experiments were performed with or without 250 μM ZnCl_2_ in two biological replicates.

However, in zinc depleted conditions, the Δ*znuCBA* mutant strain produced less biofilm than the other strains. Zinc depletion caused an overall drop in biofilm formation (Fig. 5d). These data indicated that ZnuABC is important for biofilm formation under zinc deficiency in *C. violaceum*. We wondered if zinc could be important for *C. violaceum* in the context of interspecies competition. In liquid co-culture assays with or without zinc supplementation, the Δ*znuCBA* mutant strain was outcompeted by *Burkholderia cepacia* and *S.* Typhimurium (Fig. 5e), indicating that ZnuABC is important for *C. violaceum* to compete with other bacteria.

### ZnuABC contributes to *C. violaceum* virulence and resistance to neutrophils

To verify whether zinc-related genes are involved in the pathogenesis of *C. violaceum*, we performed virulence assays in BALB/c mice, as previously determined (27, 28, 37). The Δ*zur* mutant strain was as virulent as the WT strain (most animals died 2 days post infection) (Fig. 6a). On the other hand, when infected with the Δ*znuCBA* mutant strain, all animals remained alive 10 days post infection, demonstrating that the absence of *znuCBA* renders *C. violaceum* highly attenuated for virulence. This phenotype was partially rescued in the complemented strain (Fig. 6a). We also evaluated the *C. violaceum* ability to colonize the host by assessing the bacterial CFU in mice 16 hours post infection. The livers of mice infected with the Δ*znuCBA* mutant strain had lower bacterial burdens than that of the WT or complemented strain (Fig. 6b). Additionally, we tested the survival of *C. violaceum* when incubated with neutrophils isolated from the peritoneal cavity of BALB/c mice. The CFU of the Δ*znuCBA* mutant strain decreased in the presence of neutrophils, while the CFU of the WT and complemented strains remained similar to that of these strains maintained in the control situation, without neutrophils (Fig. 6c). Altogether, these experiments showed that ZnuABC plays a key role in the virulence of *C. violaceum*, most likely by contributing to zinc acquisition in the context of infection, where neutrophils release CP to impose zinc limitation.

**Fig 6.**
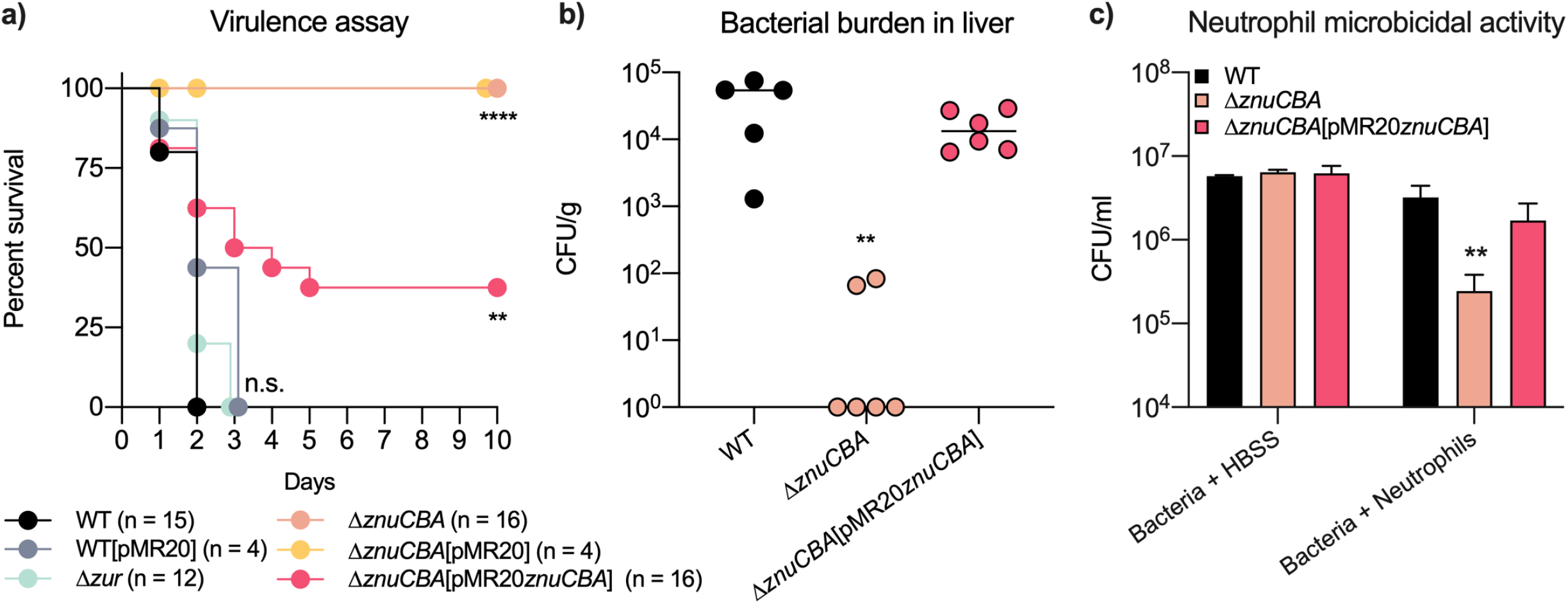
The Δ *znuCBA* mutant strain is highly attenuated for virulence in mice and susceptible to neutrophil killing. (A) Survival plot of infected BALB/c mice. Animals were i.p. injected with 10^6^ CFU of the indicated strains. Animal survival was monitored for up to 10 days. Statistical analysis was performed by the log rank (Mantel-Cox) test; \*\**P*=0.0074, \*\*\*\**P*<0.0001. (B) Bacterial burden in the liver. BALB/c mice (n=5 for WT; n=6 for Δ*znuCBA* and Δ*znuCBA*[pMR20*znuCBA*]) were infected with 10^6^ CFU. After 16 h of infection, livers were collected, homogenized in PBS, diluted, and plated for CFU quantification. Statistical analyses were performed by One-way ANOVA followed by Dunnett’s multiple-comparison test, comparing with the WT strain. \*\**P*=0.0047. (C) Neutrophil microbicidal activity. Neutrophils were infected *in vitro* with the *C. violaceum* strains at a MOI of 10 and incubated at 37°C for 2 h. The neutrophils were lysed, and diluted samples were plated on LB for bacterial CFU quantification. As a control, bacteria incubated with HBSS and maintained in the same conditions were also quantified. Statistical analyses were performed by One-way ANOVA followed by Dunnett’s multiple-comparison test, comparing with the WT strain. \*\**P*=0.0053.

## DISCUSSION

In this study, we characterized the zinc uptake transporter ZnuABC in the environmental pathogen *C. violaceum*. We described the role of ZnuABC in several zinc-dependent processes, including overcoming host nutritional immunity during infection. The *znuCBA* genes are transcribed in an operon with *zur*, and two genes encoding hypothetical proteins. This operon is repressed by Zur and derepressed in the presence of the host protein CP and the synthetic zinc chelator EDTA. We showed that the deletion of *znuCBA* renders *C. violaceum* more sensitive to zinc-depleted conditions. The compromised zinc supply affected violacein production, swimming motility, biofilm formation, and bacterial competition. Furthermore, we demonstrated, using a murine model of infection, that ZnuABC is required for *C. violaceum* virulence, liver colonization, and resistance to neutrophil.

The ZnuABC transporters have been characterized as high affinity zinc importers in several Gram-negative bacteria (3, 10, 19, 23, 24). Although our study lacks a biochemical assay to demonstrate the zinc uptake activity of ZnuABC directly, we provide evidence that ZnuABC may act as a high affinity zinc transporter in *C. violaceum*: (i) the *znuCBA* genes are part of a Zur-repressed operon that respond to zinc limitation; (ii) the absence of *znuCBA* impairs the growth of *C. violaceum* only under severe zinc scarcity, when the intracellular zinc pool was previously depleted with EDTA or CP. In several bacteria, such as *A. baumannii*, *Agrobacterium tumefaciens*, and *Pseudomonas aeruginosa*, the genes *znuA*, *znuB*, *znuC*, and *zur* are located near in the genome; commonly, *zur* is transcribed together with *znuCB* and divergently from *znuA* (24, 38, 39). In *C. violaceum*, *znuCBA* compose an operon with *zur*, CV_RS15040, and CV_RS15045. The gene CV_RS15040 encodes a small hypothetical protein (63 aa) with a predicted helical transmembrane segment. It would be interesting in future studies to address whether CV_RS15040 could be an accessory protein of the ZnuABC system. The gene CV_RS15045 encodes a putative GTPase of the COG0523 family of metallochaperones (40). Studied members of this family involved with the response to zinc limitation include YeiR of *E. coli* (41), YciC of *Agrobacterium tumefaciens* (38), ZagA of *Bacillus subtilis* (42), and ZigA of *A. baumannii* (43, 44).

Our data indicated that *C. violaceum* requires the ZnuABC system to supply zinc to several physiological processes, such as violacein production, swimming motility, biofilm formation, and bacterial competition. It has been described the requirement of zinc for enzymes of the violacein biosynthetic pathway in *C. violaceum* (45), for flagellar motility in *Proteus mirabilis*, *S.* Typhimurium, and *P. aeruginosa* (46–48), for biofilm formation via surface adhesion proteins in *Staphylococcus epidermidis* and *Staphylococcus aureus* (49), and for bacterial competition in a context of polymicrobial infections (19, 50). Although further studies are needed to understand how these processes were impaired under zinc deficiency in *C. violaceum*, our data suggest an adaptive response to ensure the zinc supply for key metabolic pathways. A global investigation of the Zur regulon could elucidate these aspects of the zinc homeostasis in *C. violaceum*.

Zinc acquisition is crucial for bacterial pathogenesis, but the contribution of the ZnuABC transporters varies according to the bacterial pathogen. The involvement of ZnuABC in virulence has been described in *S.* Typhimurium (51), *A. baumannii* (23, 24), and *E. coli* (52). Previous work from our group has indicated that *C. violaceum* relies on siderophores to overcome the iron limitation imposed by the host during infection (27, 28), but the role of zinc for the *C. violaceum* virulence remained unknown. In this study, we determined that the ABC-type transporter ZnuABC is essential for the pathogenesis of *C. violaceum* in mice, given that a Δ*znuCBA* mutant strain was highly attenuated for virulence. The low capacity of Δ*znuCBA* to colonize the liver, grow in the presence of CP, and resist neutrophil killing, indicate that ZnuABC is a primary mechanism for *C. violaceum* to overcome zinc limitation in the host.

## MATERIALS AND METHODS

### Bacterial strains, plasmids, and growth conditions

The bacterial strains and plasmids used in this work are described in Table 1. *E. coli* strains were cultivated in Luria-Bertani (LB) medium. *C. violaceum* strains were cultivated at 37°C in LB or M9 minimal medium supplemented with 0.1% casein hydrolysate (M9CH) (27, 28). Kanamycin (50 μg/ml), ampicillin (100 μg/ml) or tetracycline (10 μg/ml) were used in the cultures, when required. Conditions of zinc sufficiency were achieved by supplementation with ZnCl_2_. Zinc deficient conditions were determined by minimal inhibitory concentration (MIC) experiments, using the chelators ethylenediamine tetraacetic acid (EDTA) and N,N,N’,N’-tetrakis (2-pyridylmethyl) ethylenediamine (TPEN). For iron, we used FeSO_4_ (iron sufficiency) or 2,2’-dipyridyl (2,2’-DP) (iron deficiency), as previously defined for *C. violaceum* (27, 28).

**Table 1.** Bacterial strains and plasmids.

| Strain or plasmid | Description<sup>a</sup> | Reference or source |
| --- | --- | --- |
| *Escherichia coli* |  |  |
| DH5α | *E. coli* strain for cloning purposes | (56) |
| S17-1 | *E. coli* strain for plasmid mobilization | (57) |
| *Chromobacterium violaceum* |  |  |
| WT | *C. violaceum* ATCC 12472 WT strain with sequenced reference genome | (58) |
| WT[pMR20] | WT strain harboring the empty pMR20 plasmid | This work |
| Δ*zur* | WT strain with *zur* (CV_RS15050) deleted | This work |
| Δ*znuCBA* | WT strain with *znuC*, *znuB*, and *znuA* (CV_RS15035, CV_RS15030, and CV_RS15025) deleted | This work |
| Δ*znuCBA*[pMR20] | Δ*znuCBA* strain harboring the empty pMR20 plasmid | This work |
| Δ*znuCBA*[pMR20*znuCBA*] | Δ*znuCBA* mutant complemented with the genes *znuCBA* | This work |
| WT[p*zur-lacZ*] | WT strain with fusion of the promoter region of *zur* to *lacZ* | This work |
| Δ*zur*[p*zur-lacZ*] | Δ*zur* mutant with fusion of the promoter region of *zur* to *lacZ* | This work |
| *Burkholderia cepacia* | *B. cepacia* ATCC 17759 WT strain with sequenced reference genome | (59) |
| *Salmonella* Typhimurium | *S. Typhimurium* ATCC 14028 WT strain | (60) |
| Plasmids |  |  |
| pNPTS138 | Suicide vector containing *oriT*, *sacB*, Kan<sup>r</sup> | M. R. K. Alley |
| pMR20 | Broad-host-range low-copy vector containing *oriT*, Tet<sup>r</sup> | (61) |
| pGEM-T easy | Cloning plasmid, Amp<sup>r</sup> | Promega |
| pRK*lacZ*290 | pRK2-derived vector with promoterless *lacZ* gene, Tet<sup>r</sup> | (62) |

<sup>a</sup>Kan, kanamycin; Tet, tetracycline; Amp, ampicillin; r, resistance.

### *In silico* prediction of a Zur binding site

To find a putative Zur box in *C. violaceum*, we searched for conserved DNA binding motifs upstream of *zur*, using the MEME (Multiple Em for Motif Elicitation) tool (34). The DNA sequences (300 bp upstream of the *zur* start codon) used in the MEME analysis were obtained from 23 draft genomes of species belonging to the *Chromobacterium* genus, using the Integrated Microbial Genomes and Microbiomes (IMG/M) website (53). The DNA binding motifs predicted by MEME were searched against databases of known DNA binding motifs, using the Tomtom motif comparison tool (54).

### Construction of *C. violaceum* mutant and complemented strains

To generate the Δ*zur* and Δ*znuCBA* mutant strains, we used a mutagenesis protocol based on two-step homologous recombination and sucrose counter-selection, as previously established for *C. violaceum* (27, 28). The primers used for cloning, sequencing, and mutant confirmation are listed in Table 2. For complementation of the Δ*znuCBA* mutant strain, the three genes were PCR-amplified from the WT strain and cloned into the replicative plasmid pMR20. The primers used for cloning are listed in Table 2. The construct was mobilized into the Δ*znuCBA* mutant strain by conjugation (27, 28).

**Table 2.**
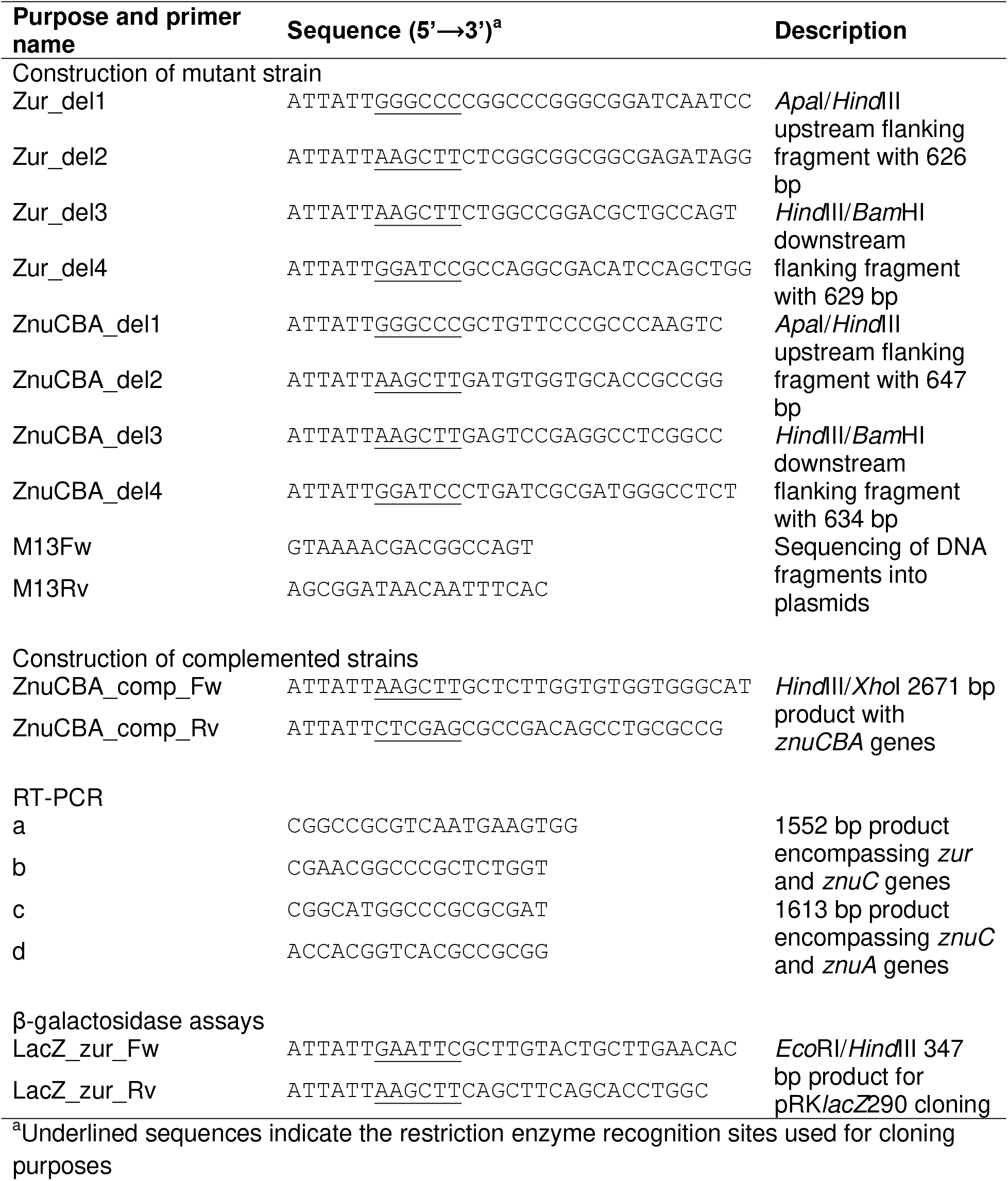
Primers used in this work.

### RNA isolation and reverse-transcriptase polymerase chain reaction (RT-PCR)

Wild-type *C. violaceum* was grown at 37°C in LB medium until the mid-exponential growth phase, and the cells were harvested by centrifugation. The total RNA was extracted using TRIzol reagent (Invitrogen) and purified with Direct-zol RNA Miniprep Plus (Zymo Research), following the manufacturer’s instructions. Purified RNA was tested in PCR reactions with the same set of primers used in RT-PCR (Table 2) to check for genomic DNA contamination. The co-transcription was analyzed by RT-PCR using SuperScript III One-Step RT-PCR System with Platinum Taq DNA Polymerase (Invitrogen), following the manufacturer’s instructions.

### Calprotectin IC_50_ assays

CP was purified as previously described (21) and stored in a CP buffer (100 mM NaCl, 3 mM CaCl_2_, 20 mM Tris pH 7.5, and 5 mM 2-mercaptoethanol). The integrity of the WT and ΔSI/SII versions of purified CP was confirmed by SDS-PAGE analysis. Two bands, one of 8 kDa (S100A8) and the other of 14 kDa (S100A9), were observed (data not shown). To assess the CP antimicrobial activity against *C. violaceum*, the 50% growth inhibitory concentration (IC_50_) of CP was calculated as previously described (17) with minor modifications. Cultures of *E. coli* DH5α and *C. violaceum* were diluted to an OD_600_ of 0.1 in LB and CP buffer (38% LB, 62% CB buffer), and WT or ΔSI/SII CP were added at several quantities (serial dilution from 960 μg/ml to 120 μg/ml). After 8 h incubation in a 96-well plate with agitation of 120 rpm, the OD_600_ was measured, and the IC_50_ was calculated in GraphPad Prism version 8 (GraphPad, San Diego, CA). The experiments were performed in three independent biological replicates.

### Construction of transcriptional *lacZ* fusions and β-galactosidase activity assay

The upstream region of *zur* (CV_RS15050) was PCR-amplified using proper primers (Table 2). The product was cloned directly into the pGEM-T easy plasmid (Promega), and subcloned into the pRK*lacZ*290 vector, as an *Eco*RI/*Hind*III fragment (Table 1). The construct containing the *zur*-*lacZ* transcriptional fusion was introduced into the *C. violaceum* strains. The strains harboring this construct were grown until the mid-exponential phase in LB medium, and were either untreated or treated with the indicated concentrations of ZnCl_2_, TPEN, EDTA, 2,2’-DP, or FeSO_4_ for 2 h. For the CP-treated cells, overnight cultures were diluted 1/6 in LB medium and CP buffer (38% LB, 62% CB buffer). Then, 500 μg/ml CP or 1 mM EDTA were added, and after 4 h the promoter activity was measured. The β-galactosidase activity assay was performed as previously described (55), using cultures from three independent biological replicates.

### Minimal Inhibitory Concentration (MIC) assay

To assess the effects of the metal chelator EDTA, MIC assays were performed with the WT and Δ*znuCBA* strains. Overnight cultures were diluted to an optical density at 600 nm (OD_600_) of 0.01 in LB medium in the presence of serially-diluted concentrations of EDTA. After 24 h cultivation at 37°C with agitation (250 rpm), the OD_600_ of all samples was measured. The experiments were performed in three independent biological replicates.

### Disk diffusion assays

To test the sensitivity to EDTA or 2,2’-DP, overnight cultures were diluted to an OD_600_ of 0.1, and 500 μl were spread on LB agar plates. Sterile paper disks were placed on the cultures. The indicated concentrations of EDTA or 2,2’-DP were dropped in each paper disk (10 μl). After 24 h incubation at 37°C, the zone of growth inhibition was measured. The experiments were performed in three independent biological replicates.

### Growth curves

Overnight cultures of *C. violaceum* were diluted in 10-15 ml of LB medium to an OD_600_ of 0.01. When necessary, the pre-inocula were treated with 1 mM EDTA, 500 μM CP WT or 500 μM CP ΔSI/SII. For the growth curves, in LB or LB plus the indicated treatments, the diluted cultures were grown at 37°C under agitation (250 rpm), and aliquots were withdrawn for OD_600_ measurement in an Eppendorf BioPhotometer. Growth curves with CP were performed in diluted LB (38% LB, 62% CB buffer). All experiments were performed in three independent biological replicates.

### Swimming motility

The swimming motility assay was performed in M9CH 0.3% agar plates as previously described (27). When indicated, the M9CH plates were supplemented with 1 mM EDTA and/or 250 μM ZnCl_2_. Swimming motility was determined as the halo diameter of bacterial spreading at 24 h and 48 h. The experiments were performed in three independent biological replicates.

### Static biofilm measurement

All strains were cultured in glass tubes without agitation in LB medium with or without 100 μM EDTA at 37°C for 24 h, before staining with 0.1% (w/v) crystal violet. Stained biofilms were resuspended with 100% ethanol (1 ml), followed by measurement of OD_600_ (27). The experiments were performed in six biological replicates. Data are plotted as the ratio of biofilm measurement by bacterial growth.

### Interspecies competition

The *C. violaceum* WT and Δ*znuCBA* strains were co-cultivated with other bacteria. The *C. violaceum* strains (resistant to ampicillin) and the ampicillin-sensitive bacteria *B. cepacia* and *S.* Thypimurium were grown overnight in LB. Bacterial cultures were diluted 1:20 in LB or LB plus 250 μM ZnCl_2_ and mixed 1:1. After incubation for 2 h at 37°C and 250 rpm, the co-cultures were serially diluted and plated in selective LB ampicillin plates for CFU enumeration. For comparison, the CFU of each bacterium grown in the same conditions, but without co-cultivation, was also determined in LB plates.

### Mouse virulence assays

6-week-old female BALB/c mice were used to inject intraperitoneally 10^6^ CFU from the *C. violaceum* strains, as previously determined (27, 28, 31, 37). Animal survival was followed up for 10 days post infection. To determine the bacterial burdens in the liver, mice were infected as previously stated and euthanized after 16 h. Livers were aseptically collected and blended in PBS. Serial dilutions were plated on LB for CFU quantification. For the *in vitro* neutrophil killing assays, we used a previously described protocol of thioglycolate-induced peritonitis to obtain a neutrophil enriched fraction from the peritoneal cavity of BALB/c mice (28). The *C. violaceum* strains were added to the neutrophils in a Hank’s Balanced Salt Solution (HBSS) supplemented with 2% fetal bovine serum at a multiplicity of infection (MOI) of 10:1. After 2 h incubation at 37°C with gentle agitation, the neutrophils were lysed by adding an equal volume of water. The vortexed suspensions were serially diluted and plated in LB for CFU quantification. The CFU of the strains grown in HBSS was determined, as a control. The procedures with mice were performed under the Ethical Principles in Animal Research adopted by the National Council for the Control of Animal Experimentation (CONCEA). The protocol (181/2017) was approved by the Local Ethical Animal Committee (CEUA) of FMRP-USP.

### Statistical analysis

Statistical analysis was performed using GraphPad Prism version 8 (GraphPad, San Diego, CA). Data distribution was assessed using a Shapiro-Wilk test of normality, assuming statistical significance at 0.05. Multiple comparison tests were performed and are indicated in each figure.

## ACKNOWLEDGMENTS

This research was supported by grants from the São Paulo Research Foundation (FAPESP; grants 2018/01388-6 and 2020/00259-8) and Fundação de Apoio ao Ensino, Pesquisa e Assistência do Hospital das Clínicas da FMRP-USP (FAEPA). Calprotectin research in the Skaar and Chazin laboratories is supported by grants R01 AI101171 and R01 RAI150701 from the US National Institutes of Health. During the course of this work, RERSS (grant 2017/03342-0) and WPSJ (grant 2018/14737-9) were supported by FAPESP fellowships.

